# Quantitative, super-resolution localization of small RNAs with sRNA-PAINT

**DOI:** 10.1101/716696

**Authors:** Kun Huang, Feray Demirci, Mona Batish, Blake C. Meyers, Jeffrey L. Caplan

## Abstract

Small RNAs are non-coding RNAs that play important roles in the lives of both animals and plants. They are 21- to 24-nt in length and around 10 nanometers (nm) in size. Their small size and high diversity have made it challenging to develop detection methods that have sufficient resolution and specificity to multiplex and quantify. We created a method, sRNA-PAINT, for the detection of small RNAs with 20 nm resolution by combining the super-resolution method, DNA-based points accumulation in nanoscale topography (DNA-PAINT), and the specificity of locked nucleic acid (LNA) probes for the *in situ* detection of multiple small RNAs. The method relies on designing probes to target small RNAs that combine DNA oligonucleotides (oligos) for PAINT with LNA-containing oligos for hybridization; therefore, we developed an online tool called “Vetting & Analysis of RNA for *in situ* Hybridization probes” (VARNISH) for probe design. Our method utilizes advances in DNA-PAINT methodologies, including qPAINT for quantification, and Exchange-PAINT for multiplexing. We demonstrated these capabilities of sRNA-PAINT by detecting and quantifying small RNAs in different cell layers of early developmental stage maize anthers that are important for male sexual reproduction.

## Introduction

In plants, 21- to 24-nucleotide (nt), non-coding small RNAs (sRNAs) regulate many important biological processes (1). Plant and animal genomes encode various sRNAs that can be divided into two major categories: microRNAs (miRNAs) and small interfering RNAs (siRNAs) (2). miRNAs are derived from long, hairpin precursor RNAs processed by precise cleavage, yielding mature miRNAs which influence gene transcript levels and translation (3). Plant siRNAs, also derived from long precursors that are made double-stranded by RNA-dependent RNA polymerases (RDRPs), are processed into mature siRNAs, and regulate gene transcript levels through post-transcriptional silencing, and epigenetically via RNA-directed DNA methylation (RdDM) (4). Some plant miRNAs trigger phased, secondary siRNAs (phasiRNAs), which are either 21 or 24 nt in length (5-7). Anthers, the plant male reproductive organ, are a particularly rich source of miRNAs and phasiRNAs, well-described in grasses including maize (7); during the meiotic stage, anthers are composed of cell layers that include the epidermis, endothecium, middle, and tapetum, that all surround the pollen mother cells. In maize anthers, the abundance and distribution of miRNAs and phasiRNAs vary in different cell layers (7), and our interest in the spatial organization and concentration of these sRNAs, particularly at the subcellular level, provided us a strong motivation to develop new methods for fluorescent detection.

sRNAs are mobile, moving within and between organelles, from cell-to-cell, and long distances within an organism or even across organismal boundaries in plant-pathogen interactions (8). For example, within a cell, sRNAs move between the nucleus and cytoplasm, bind Argonaute (AGO) proteins, Dicing bodies (D-bodies), Cajal bodies (CBs), and may be polysome-associated or found with organelles such as the endoplasmic reticulum (9-12). Intercellular movement of sRNAs, presumably via plasmodesmata in plants, regulates tissue and organ development by silencing and regulating developmentally important genes (13). Systemic translocation of sRNAs has been demonstrated by epigenetic silencing across graft junctions and sRNA profiling of vasculature (8, 14). Remarkably, sRNAs also move between organisms at host-pathogen interfaces (15). Pathogens, including parasitic plants, regulate host defense responses via miRNAs and *trans*-species sRNAs, and vice versa (16). In animals, extracellular vesicles (EVs) transport miRNAs from cell-to-cell and over long-distances (17). Plant EVs are populated with miRNAs as well as with fragments as small as 10 to 17 nt, so-called tiny RNAs (2). All of these sRNAs vary in abundance from low to extremely high, as measured by sequencing, which typically utilizes gram quantities of plant tissue, a process that substantially limits spatial analyses. At the subcellular level, the localizations of these RNAs are not well-characterized. Thus, future approaches to sRNA characterization will require nanometer resolution, high specificity, and both multiplexing and quantitative capabilities.

For *in situ* hybridization of sRNAs, the use of locked nucleic acid (LNA) oligonucleotide probes has shown specificity and high fidelity (18). However, colorimetric and even fluorescent localization of sRNAs using LNA-based methods are able to resolve only cell-level localization patterns. Because of the limits of light diffraction, light microscopy cannot resolve spots below ∼200 nm in diameter and 400 to 700 nm in axial length (19). New techniques have been developed to drive microscopic detection beyond the diffraction limit, yielding so-called “super-resolution” microscopy, such as Photo-Activated Localization Microscopy (PALM), Structured Illumination Microscopy (SIM), direct Stochastic Optical Reconstruction Microscopy (dSTORM), and Points Accumulation for Imaging in Nanoscale Topography (PAINT) (19). We have previously demonstrated an sRNA-FISH method combined with SIM and dSTORM (20). However, dSTORM utilizes the blinking property of fluorescent dyes and relies on specific excitation and buffer conditions to achieve proper imaging (21). Photoswitching of the dye molecule is hard to predict, which made absolute quantification difficult usingthe dSTORM method. The number of dyes with the appropriate photoswitching properties within the visible spectrum is also a limiting factor for multiplexed detection (21). DNA-PAINT decouples the blinking events from dye photophysics, using the binding and dwelling kinetics of a short dye-labeled oligonucleotide “imager strand” to localize target-specific “docking strands” (22, 23). The docking strand in DNA-PAINT is linked to an antibody for single-molecule protein detection, with imager strands introduced by perfusion, one imager strand at a time, each specific to a given docking strand and protein. After the image is acquired, the imager strand is washed off and then the next imager strand is introduced (21). As a result, a single dye can be linked to different imager strands and applied for detection of numerous protein targets via use of different imager strand-docking strand combinations. This multiplexing technique, known as “Exchange-PAINT” has been used for *in situ* imaging of protein targets, such as microtubules and mitochondria (23), and *in vitro* profiling of miRNAs using barcoded synthetic DNA origami (24).

To achieve subcellular, nanometer resolution imaging for the localization and analysis of sRNAs, we created a detection method called sRNA-PAINT. This method combines the high resolution and precise quantification of DNA-PAINT with the efficiency and specificity of LNA-based *in situ* hybridization. We created a probe design tool for our method and show that these sRNA-PAINT probes can used for robust sRNA detection. And, we have demonstrated that it can be combined with qPAINT to localize and quantify distinct sRNAs in fixed biological sample, and with Exchange-PAINT for multiplexed target detection.

## Materials and Methods

### Plant materials and oligonucleotides

Maize anthers from W23 were provided by the Walbot lab at Stanford University (Palo Alto, CA). Plants were grown in Palo Alto, CA under greenhouse conditions. Anther dissection and measurements were performed as previously described (25). LNA-modified oligonucleotide probes were designed using our VARNISH software and synthesized by Exiqon (QIAGEN; Germantown, MD). Imager strands coupled with Alexa Fluor 647 (AF647) were ordered from IDT (Coralville, Iowa).

### Sample preparation

Sample preparation for *in situ* hybridization was performed as previously described (20). Briefly, anthers were dissected and fixed in a 50 ml tube using 4% paraformaldehyde in 1xPHEM buffer (5 mM HEPES, 60 mM PIPES, 10 mM EGTA, 2 mM MgSO_4_ at pH 7). Samples were then processed in a vacuum chamber (0.08 MPa) three times, 15 min each. After fixation, samples were embedded in paraffin at the Histochemistry and Tissue Processing Core Lab at Nemours/Alfred I. duPont Hospital for Children (Wilmington, DE). Paraffin samples were sectioned at 6 µm thickness using a paraffin microtome and dried on a Wide Spectral Band 600+/- 100 nm Gold Fiducials coverglass (600-100AuF; Hestzig LLC, Leesburg, VA) at 37 °C degree on a slider warmer.

### In situ hybridization

The *in situ* hybridization step was performed following our previously-published protocol (20). Briefly, samples were de-paraffinized using Histo-Clear (item 50-899-90147; Fisher Scientific, Pittsburgh, PA) and re-hydrated by going through an ethanol series of 95, 80, 70, 50, 30, 10% (vol/vol) and water (1 min) at room temperature. Then samples were treated with protease (item P5147; Sigma-Aldrich, St. Louis, MO) for 20 min at 37°C. Excess formaldehyde background was removed by treating samples with 0.2% glycine (item G8898; Sigma-Aldrich) for 20 min. After two washes in 1xPBS buffer (phosphate-buffered saline), samples were dehydrated by going through an ethanol series of 10, 30, 50, 70, 80, 95%, and 100% (vol/vol). Hybridization was done with 10 µM probe at 53.3°C in a hybridization oven. After hybridization, samples were washed twice with 0.2x SSC buffer (saline-sodium citrate). To immobilize the hybridized probes, samples were incubated for 10 minutes in freshly prepared EDC solution containing 0.13 M 1-methylimidazole, 300 nM NaCl (pH 8.0). Then samples were incubated for 1 hour and 15 minutes in 0.16 M N-(3-Dimethylaminopropyl)-N’-ethylcarbodiimide hydrochloride (EDC) (item 03450, Sigma-Aldrich, St. Louis, MO) solution. Slides then were washed twice in TBS solution, 10 min each wash. Hybridized samples were kept in 1xTBS at 4°C until imaging. smFISH was carried out as previously described (26). 35 probes were used to detect the *PHAS* precursor lncRNA, as shown in Supplementary Table 2. Quantification of smFISH was processed using SpotCounter program in ImageJ. Three different spots from each layer, were used for quantification, which were the exact same locations used in qPAINT quantification for direct comparison.

### Super-resolution imaging and image reconstruction

Super-resolution imaging was carried out on an inverted Zeiss Elyra PS.1 super-resolution microscope (Carl Zeiss, Lberkochen, Germany). TIRF illumination was done using a 100% 642 nm laser and α-Plan-Apochromat 100x/1.46 oil objective. For each imager strand, as well as control imager strand, images were taken with an exposure time of 100 ms, an EMCCD Gain 30, and 20,000 frames in total.

Each image was analyzed and rendered in Zen software (Carl Zeiss Inc., Thornwood, NY). The images were processed using the following identical parameters: Ignore overlapping molecules; Peak Mask Size (6); Peak Intensity to Noise (6.0); Fit model (x,y 2D Gauss Fit). Drift correction was done by using fiducial-based algorithm. For image rendering, pixel resolution was set to 16 nm/pixel and 2 nm/pixel for zoomed in images with 1x and 0.5x PSF (point spread function) expansion factor. The number of photons was selected between 420 and 10000 to eliminate non-specific background.

### sRNA-PAINT

A DH40iL culture dish incubate system (model 640388; Warner Instruments LLC, Hamden, CT) and a quick release magnetic chamber for 25 mm low profile, round coverslips (model 641943; Warner Instruments LLC) were assembled and used as the perfusion chamber. A ValveLink8.2 Perfusion System (AutoMate Scientific, Berkeley, CA) was used for perfusing buffer and imager strand solutions and washing solution into the chamber. ValveLink 8.2 perfusion system is a gravity force-driven perfusion system. We used a working height about 15 cm above the imaging chamber. All imager strands were diluted to 0.5-2 nM in buffer C (1× PBS, 500 mM NaCl, pH 8). Images were taken with constant flow of imager stand. A Masterflex C/L peristaltic pump (60 RPM, model 77120-62; Cole-Parmer Instrument Company LLC, Vernon Hills, IL) was used to constantly collect the waste buffer flow at maximum speed.

### sRNA-Exchange-PAINT

For multiplexed detection, all the probes were designed with VARNISH tool by picking different docking and imager strand combinations. Following the same *in situ* hybridization procedure as described earlier, each probe was denatured in an individual tube. After chilling on ice, all the probes were mixed together and hybridized to the tissue simultaneously overnight at 53.3°C in a hybridization oven. The probes were washed off with 0.2x SSC buffer the next morning, and stored in 1x PBS buffer till imaging. DAPI (4’,6-diamidino-2-phenylindole) stained nuclei images were taken after Exchange-PAINT. Alignment of channels were done using TrakEM2 package of ImageJ (27). Probes used for multiplexed detection are listed in Supplementary Table 1.

### Quantification of sRNA-qPAINT with Picasso software

qPAINT data analysis was performed following the protocol by Schnitzbauer et al. (21) and using the Picasso software package. In brief, 8000 frames of the raw movie file of PAINT data were processed with “Picasso: Localize”. We adjusted the threshold until only the PAINT spots were detected and selected (13000 Min Net Gradient was chosen for this manuscript). After “Localize (Identify & Fit)”, a .hdf5 file was generated and this was used as the input for the next module. The .hdf5 file was opened in “Picasso: Render”, and marker-based drift correction or redundant cross-correlative drift correction was performed. We picked a 3.18 µm (20 pixel) diameter circle in the cytoplasm of each anther cell layer using “Tools: Pick” and then performed qPAINT analysis with Picasso. The copy number in these picked regions of each cell in each layer was used to calculate the number of binding sites in 63.25 µm^2^, which is roughly the area of the cytoplasm of cells in our sections. The influx rate was calculated using the formula, *ξ* = *k*_on_ × *c*, in which *k*_on_ represented the association constant of the imager strand (1.5 * 10^6^ (Ms)^−1^), and *c* represented the concentration of the imager strand. The dark time under “View: Show info” and the calculated influx rate were used to calculate binding sites. A total of 150 sample areas across three biological replicates were used to determine the background binding number. We observed 5.12 background binding sites for scrambled control LNA probe. As a result, the 5.12 background binding sites were subtracted from all other qPAINT quantifications. For each sample, 10 locations for each cell layer were used to calculate the binding sites.

### Small RNA library and data handling

The details of the library are as follows: it is a maize small RNA library with GEO accession number GSM1262527 that includes 24,145,201 genome-matched reads between the sizes of 18 and 34 nt. After removing adapters and low-quality reads, small RNA reads length between 18 and 34 nt were mapped back to the reference genome of maize, version AGPv4 (28). Abundances of small RNAs were normalized to “TP10M” (transcripts per 10 million) based on the total count of genome-matched reads in the library.

### sRNA sequences

All small RNA probe sequences and imager strands used are listed in Supplementary Table 1.

## Results

### sRNA-PAINT probe design

Most eukaryotic sRNA sequences are between 21 to 24 nt in length. To perform sRNA-PAINT, we designed a probe that is composed of three sequences: the probe backbone sequence with LNA bases, the DNA-PAINT docking strand sequence, and a linker sequence that connects those two (Figure 1). The probe design is the most critical step of the sRNA-PAINT method; to facilitate this process, we created an online tool called VARNISH (Vetting & Analysis of RNA for *in situ* Hybridization probes) for automated design of sRNA-PAINT probes (https://wasabi.ddpsc.org/~apps/varnish/). The tool requires the input of a target sRNA sequence and the hybridization parameters (defaults provided), including sodium (50 mM), magnesium (0 mM) and temperature (25°C). VARNISH first will reverse complement the sRNA sequence and will then choose between 19 and 22 nt of the sequence such that the melting point temperature (*T*_*m*_) is lower than 60°C, with preference for lower *T*_*m*_ values for the longest sequence length. The melting temperature is calculated by using the “Analyze” function from the IDT web application program interface (API) (https://www.idtdna.com/AnalyzerService/AnalyzerService.asmx). The final sequence is the probe backbone (Figure 1, cyan). Next, LNA bases and a linker sequence are introduced to the selected probe backbone by VARNISH. The default linker sequence is “tattcgt”, which has no match to the sRNA sequences in miRBase, but which can be changed to any desired custom linker sequence. VARNISH introduces between five and nine LNA bases (default is eight) and it avoids stretches of more than four LNA bases and three or more Gs or Cs. The algorithm exhausts all the possible combinations of the positions for LNA bases, and it calculates the approximate *T*_*m*_ using previously estimated *T*_*m*_ values for different LNAs (29). Next, VARNISH takes the top 200 probe backbone candidates with the highest *T*_*m*_ and calculates the exact *T*_*m*_ using the IDT web API. The last step is to add the DNA-PAINT docking strand. In the VARNISH tool, the docking strand can be either chosen from a drop down menu containing the 13 previously used docking strands (23) or the sequence can be entered manually in the VARNISH tool. The software will conduct a homo-dimer and secondary structure analysis from all, or a subset of, the 13 provided docking strands in combination with the linker and the top 200 probe backbone candidates. These two steps are performed by using the “SelfDimer” and “UNAFoldRun” functions from the IDT web API. Finally, the algorithm will choose the top 10 candidates for each docking stand with the highest *T*_*m*_ and lowest ΔG for homo-dimer and secondary structure predictions. Once the computations are complete, VARNISH will send an email message containing a weblink to an output page that lists the 10 top probe candidates and information on ordering both the probes and their corresponding imager strands.

**Figure 1.**
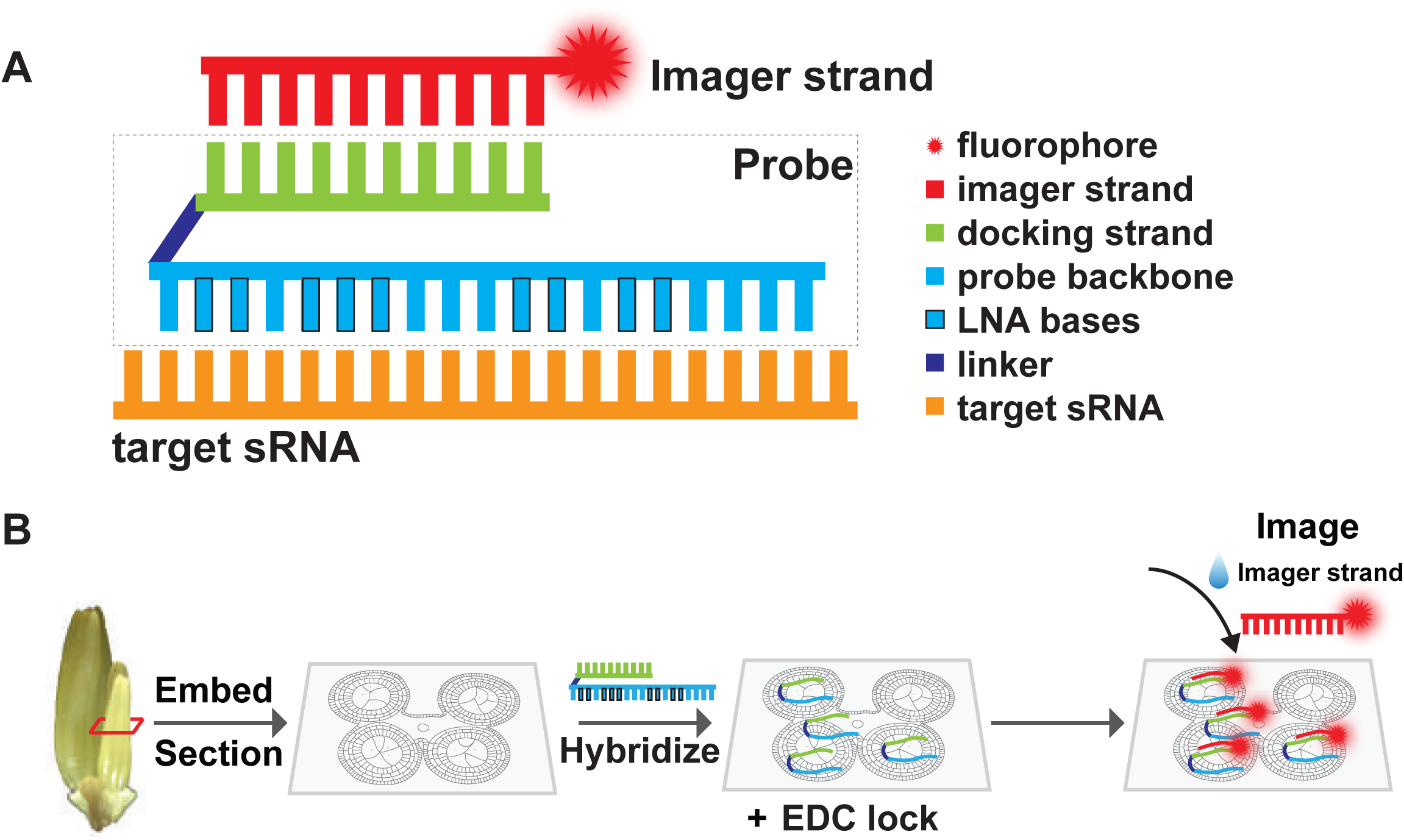
Illustration of sRNA-PAINT probe design and workflow. (**A**) Illustration of a VARNISH probe design. A VARNISH probe is comprised of three parts: a probe backbone (cyan) that is complementary to the target sRNA (orange), a docking strand (green) and a linker sequence (dark blue) connecting them. The imager strand is conjugated to the fluorophore, and it will find its corresponding docking strand during the imaging process. (**B**) A workflow for sRNA-PAINT. Samples were fixed and embedded, and a thin section is placed on the coverglass. During the hybridization process, the VARNISH probes will bind to sRNAs in the sample and locked with EDC. During the imaging step, imager strands are added to the probe-hybridized sample on the coverglass.

### sRNA-PAINT detects and quantifies cellular sRNAs at single molecule resolution

Figure 1B shows the general process for the sRNA-PAINT method. First, the samples are fixed, embedded in paraffin, and sectioned. Then, thin sections were placed onto coverslips with gold fiducials sealed with silicon dioxide (SiO_2_). The gold fiducials were used as alignment landmarks for image registration over time and after buffer exchanges. We highly recommend that no additional coatings are added, such as poly-l-lysine, since they resulted in non-specific binding. Next, during the hybridization process, probes are applied to fixed and sectioned samples (Figure 1B). Excess probes are removed in the washing process. Afterward, imager strands are perfused in. When the un-annealed imager strands flow within the buffer, the speed of the movement is too fast to be caught by the microscope. However, when an imager strand finds its docking partner, and dwells on it, the short time period before the imager strand dissociates from the docking creates a blinking event that can be detected using a total internal reflection fluorescence microscope (TIRFM). As a result, when thousands of these blink events are collected, detection of all the docking sites within the sample is eventually completed, defining the locations of the sRNA targets.

For sRNA-PAINT probe labeling, we selected Alexa Fluor 647 dye (AF647). First, AF647 emits far-red fluorescence that is distinct from tissue autofluorescence; second, its photophysical property made it a dye suitable for generating quality super-resolution images (30). The sRNA probes designed by VARNISH were hybridized and the coverslip was mounted in a perfusion chamber. The corresponding, 3’-dye-labeled imager strands were continuously perfused and imaging was conducted using TIRFM. As a demonstration, we performed sRNA-PAINT on an abundant 24-nt phasiRNA – a class of plant secondary siRNAs abundant in the early developmental stages of maize anthers (7) (Figure 2); the 24-nt phasiRNAs are triggered by an miRNA, miR2275, providing us with two different classes of small RNAs in one tissue. Our sRNA-PAINT experiment (Figure 2A) was consistent with our previous findings that the 24-nt phasiRNA was present in all of the anther cell layers, and most abundant in the tapetum layer and pollen mother cells (7) ; however, our, sRNA-PAINT approach achieved a much higher resolution of below 10 nm (Figure 2B).

**Figure 2.**
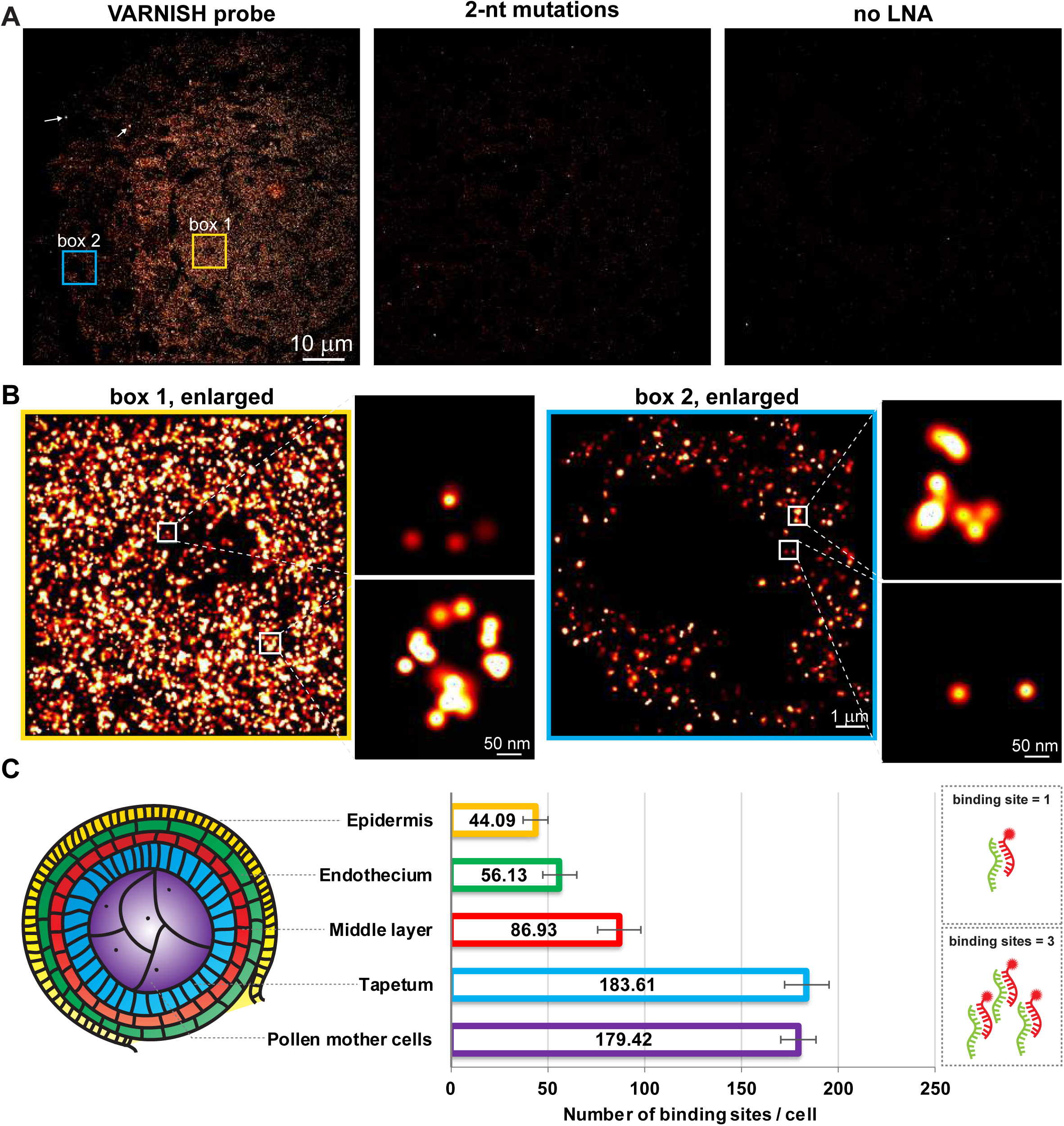
sRNA-PAINT detects 24 nt-phasiRNA specifically in an early-stage maize anther. (**A**) Detection of a 24-nt phasiRNA in a cross-section of maize anther. The VARNISH probe and modified probes with either 2-nt mutations or the LNAs were applied to samples. 24-nt phasiRNA probes showed signal while the mutation probe and no-LNA probe detected little to no signal. Scale bar = 10 µm in all images. A red-to-white lookup table was used to display low-to-high intensity values. Arrows indicate 2 out of ∼66 fiducials used in experiments. (**B**) Zoomed images of boxes 1 and 2 in panel A. There are more 24-nt phasiRNA localization events in the pollen mother cells (box 1) compared to the epidermal layer (box 2). Expanded boxed areas (white) show representative areas at higher magnification, with centroid positions shown in blue crosses. (**C**) qPAINT analysis of 24-nt phasiRNA in different anther cell layers. Cartoon diagram representing each cell layer of a cross section of a maize anther lobe: epidermis (yellow), endothecium (green), middle layer (red), tapetum (blue), and pollen mother cells (purple). Numbers shows the binding events for each cell in all cytoplasmic areas of different cell layers. 10 different locations in each specified cell layer of three biological replicates were taken for statistical analysis (n=150). Error bar indicates standard error for each sample.

qPAINT can count molecules by analyzing the predictable and programmable binding kinetics of the imager strand to the docking strand (22). This quantitative method is based solely on binding kinetics, and therefore, individual molecules do not need to be resolved. The Picasso software is publicly available and can perform this calculation for the binding sites within a selected region (21). We then performed qPAINT analysis on 24-nt phasiRNA with Picasso. As shown in Figure 2C, we detected 71.51 copies of 24-nt phasiRNA per selected region per cell in epidermal cells, 89.62 copies in endothecium cells, 131.92 copies in middle layer cells, 199.92 copies in pollen mother cells, and most abundant, 209.86 copies in tapetal cells. These copy number values are minus the background detected in the scrambled LNA control probe that was calculated to be 5.12 binding sites using 150 regions of images from three biological replicates (Supplementary Figure 1). The scrambled LNA probe used to estimate the non-specific binding of LNA probes has no similarity with maize small RNAs. Pollen mother cells and tapetum have a greater abundance of the 24-nt phasiRNA compared with the other cell layers. qPAINT, in combination with sRNA-PAINT, enabled the detection of the copy number of sRNAs in cell sections and the sRNA targets in the different cell layers.

Next, we tested the specificity of the VARNISH probes and the essentiality of LNA bases. First, we tested the specificity of the probe backbone by mutating 2 nucleotides in the 24-nt phasiRNA probe backbone. As shown in Figure 2 and quantified in Supplementary Figure 2, the probe with the 2-nt mutation detected very low background signal compared with the original probe, resulting in an average of 95% decrease of the detected spots. Second, we confirmed that LNA is essential for efficient detection of small RNA targets (18). We found that without LNA, the signal intensity is much lower compared with probes with LNA (Figure 2) and nearly a 92% reduction in binding sites quantified by qPAINT (Supplementary Figure 2). The scrambled, control LNA probe detected very little background fluorescence (Supplementary Figure 1).

After the first round of imaging, buffer was perfused into the imaging chamber, and the signal was diminished rapidly within 1 min, suggesting that stripping off the imager strand is very efficient. Next, sRNA-PAINT signal was re-achieved by re-applying the imager strands (Figure 3A). We used EDC (1-ethyl-1-(3-dimethylaminopropyl) carbodiimide) prior to imaging in order to cross-link and immobilize the probes to prevent probe stripping during wash steps (31). After EDC treatment, the signal intensity in the re-imaged sample was comparable with the original image (Figure 3A) and there was no significant difference in quantified the number of binding sites (Figure 3B). Without EDC, there was a decrease in image intensity after washing, as shown in Figure 3A and quantified in Figure 3B. The number of localization spots detected remain relatively stable during the imaging process of 20,000 frames for samples with or without EDC treatment (Supplementary Figure 3). We also confirmed that a different imager strand had no or very little non-specific binding (Figure 3A), which is similar to the original DNA-PAINT method (23).

**Figure 3.**
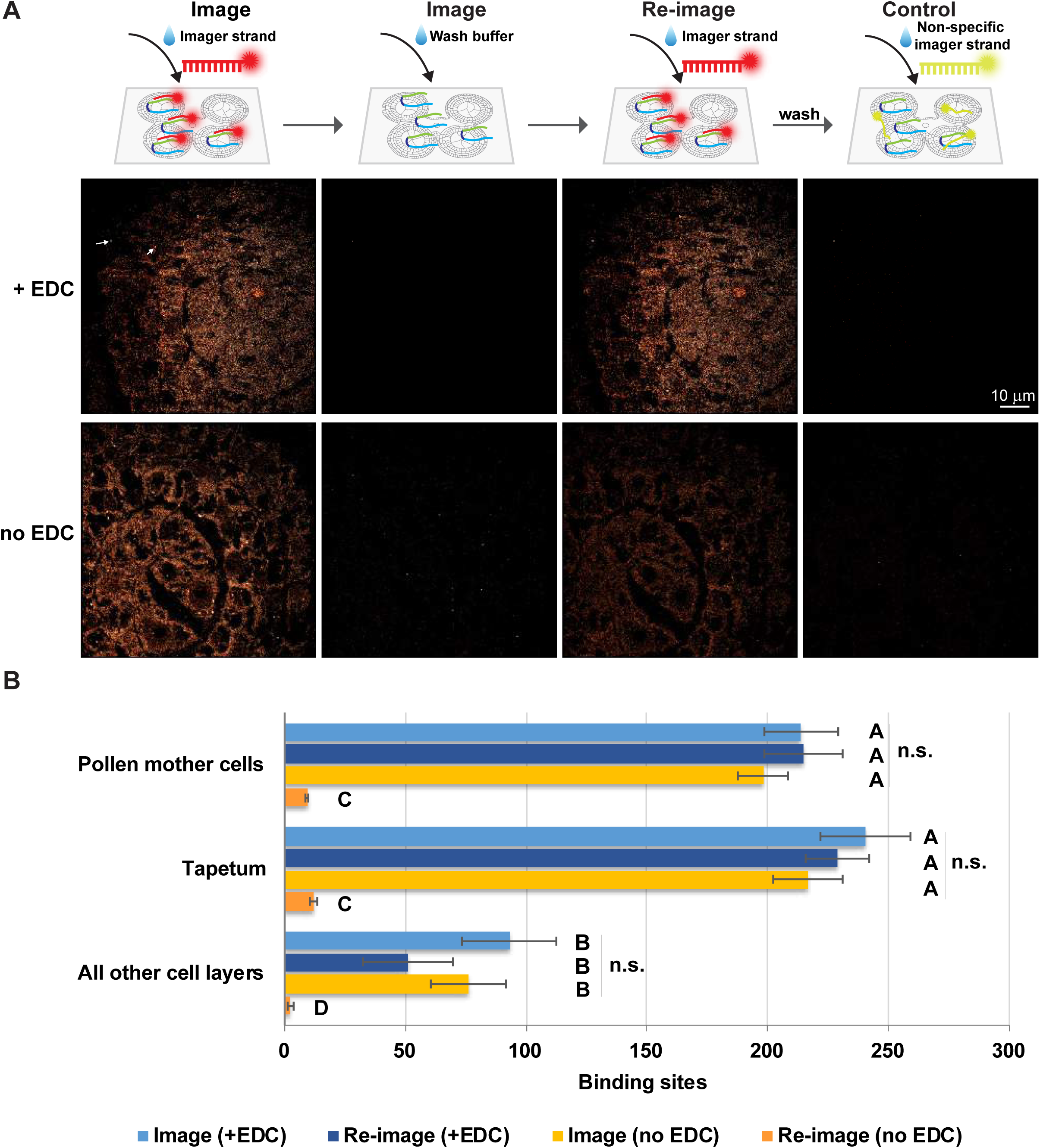
Comparison of sRNA-PAINT imaging and re-imaging with and without EDC treatment. (**A**) sRNA-PAINT imaging signal can be re-detected to a similar level when treated with EDC. Without EDC treatment, re-imaged samples have lower fluorescence signal. A non-specific imager strand (last column) was used as a control. Scale bar = 10 µm in all images. We used the same anther section used in Figure 2. Arrows indicate 2 of the of ∼66 fiducials used in experiments. (**B**) qPAINT analysis of image in panel A. Numbers shows the binding events for each cell in different cell layers. 10 different locations in each specified cell layer were taken for statistical analysis. Error bars indicate standard error for each sample. Letters indicate significant differences using Tukey’s test (P < 0.05, n=50). No significant difference was observed in each cell layer during imaging and re-imaging in EDC treated samples.

### Sequential multiplexed imaging of phasiRNA biogenesis components using sRNA-Exchange-PAINT

Sequential detection of sRNAs can be performed by multiplexed VARNISH probes with different docking strand combinations. We name this technique sRNA-Exchange-PAINT, as a variation of the protein-focused, multiplexed Exchange-PAINT method. For this analysis, we measured both miR2275 and the most abundant 24-nt phasiRNA it triggers, in maize anthers. As shown in Supplementary Figure 4, the imager strand used for miR2275 (P0*) and 24-nt phasiRNA (P1*) do not non-specifically bind to each other’s docking strand. Sections on coverglass were mounted into a perfusion chamber and were first perfused with P0* imager strand. After the image of miR2275 was acquired, buffer wash was performed to completely remove the P0* imager strand. Next, the imager strand P1* was applied, and generated the image for the 24-nt phasiRNA. Using traditional *in situ* hybridization methods, miR2275 and 24-nt phasiRNA were thought to co-localize in the tapetum layer and pollen mother cells (7). Using sRNA-PAINT, which has a resolution down to 10 nm, we confirmed that they co-localize to the tapetum layer and pollen mother cells within the anther tissue, but we were not able to detect significant co-localization at the level of single molecule analysis (Supplementary Figure 4B, zoomed image).

Higher levels of multiplexing can be achieved by sequential detection of multiple small RNA targets. To achieve this, different docking strands could be attached to different sRNA-PAINT probe backbones by simply specifying the choice of docking strand in the VARNISH software. All the sRNA-PAINT probes are then hybridized to the tissue together in one hybridization step; after stringent washes, each imager strand is perfused in for sequential detection of each target. We tested this high-level multiplexing with the phasiRNA biogenesis components in 1 mm premeiotic maize anthers (Figure 5A). We aimed for four components of the phasiRNA biogenesis pathways that shared the same precursor: miR2275, the *PHAS* lncRNA precursor, and two phasiRNAs (5 phase “cycles” apart, or 5 × 24 nt = 120 nt apart). We also included a fifth candidate, miR166, which regulates flower development but does not belong to the 24-nt phasiRNA biogenesis pathway (32). Each of the five sRNA-PAINT probes, used at the same concentration, was hybridized simultaneously to a maize anther cross-section (Figure 5A). We observed effective detection of the targets by exchanging imager strands (Figure 5B); DAPI was subsequently imaged as a counterstain for nuclei. In comparison with sRNA-seq data of multiple pooled 1 mm anthers, quantification of each sRNA-PAINT channel showed similar trends in sRNA abundances (Figure 5C). Single-molecule resolution was preserved with high-level multiplexing (Figure 5D zoom) demonstrating that sRNA-Exchange-PAINT preserved the quantification capability of qPAINT.

**Figure 4.**
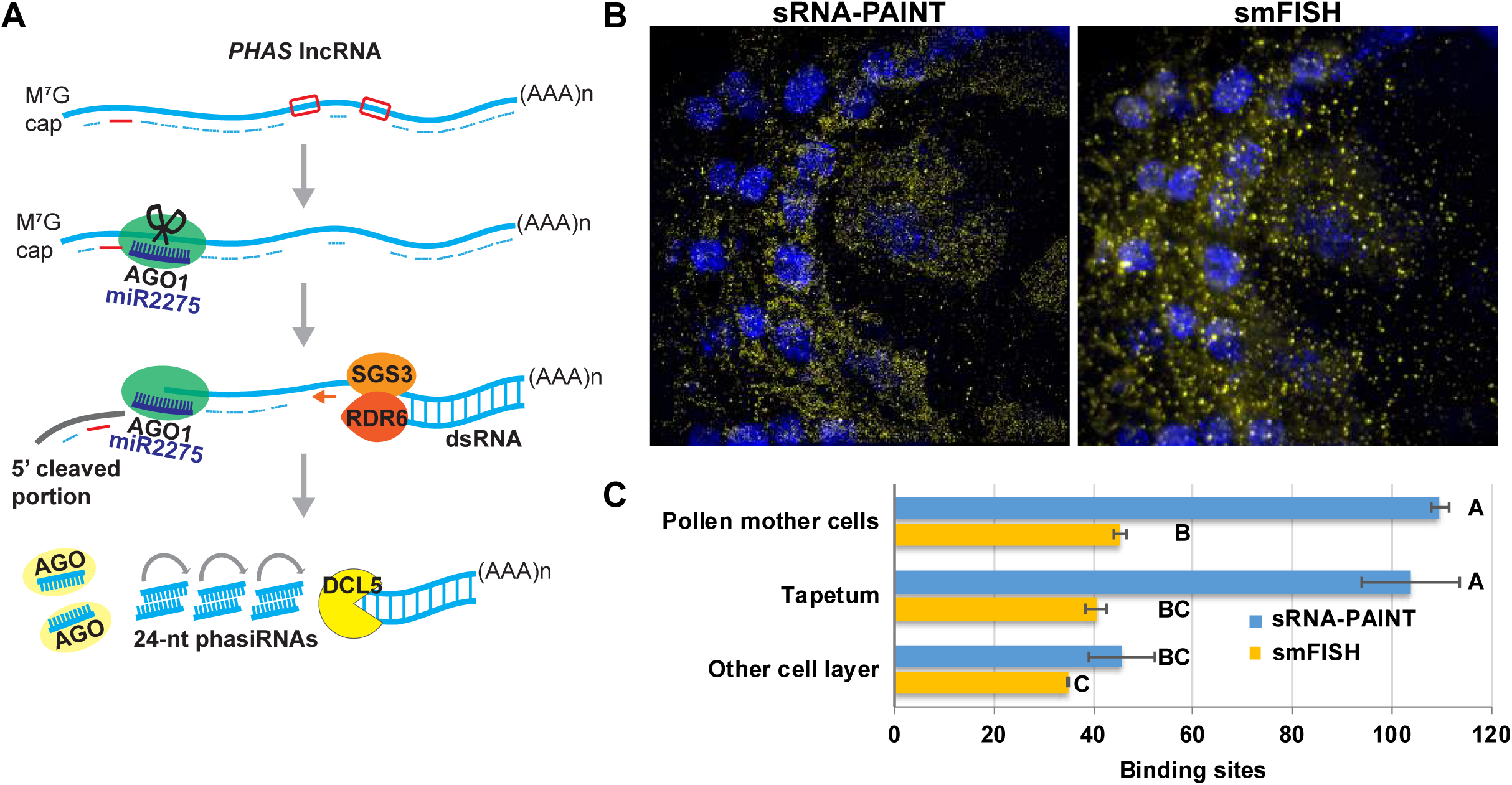
Application of sRNA-PAINT for detecting lncRNA transcript. (**A**) biogenesis pathway of phasiRNA in maize. AGO1-miR2275 cleaved *PHAS* lncRNA is produced into long dsRNA via RDR6-SGS3. DCL5 cleaves target sites into 24-mers (phasiRNAs). Our sRNA-PAINT suggests that AGO1-miR2275 and RDR6-SGS3 might be present simultaneously on the *PHAS* lncRNA. The AGO1-miR2275 cleaved 5’ unpaired portion may not go through immediate degradation. Red bar represents the relative location of the VARNISH probe for *PHAS* lncRNA. Red boxes represent probe locations for 24-nt phasiRNA 1 and 24-nt phasiRNA 2. smFISH probes are represented with dash lines. (**B**) Comparison of sRNA-PAINT and smFISH for the same RNA transcript (*PHAS* lncRNA). sRNA-PAINT detected a similar RNA localization pattern compared with smFISH, with increased single-molecule resolution and a higher quantification count. All images were taken on the same coverslip sequentially. Scale bar = 10 µm in all images. (**C**) Quantification analysis of image in panel A. sRNA-PAINT detected more *PHAS* lncRNA than smFISH. Numbers show the binding events for each cell (90-pixel size diameter circle in Picasso software) in different cell layers. 3 different locations in each specified cell layer were taken for statistical analysis. Different letters indicate significant differences (Tukey’s test; P < 0.05, n=90).

**Figure 5.**
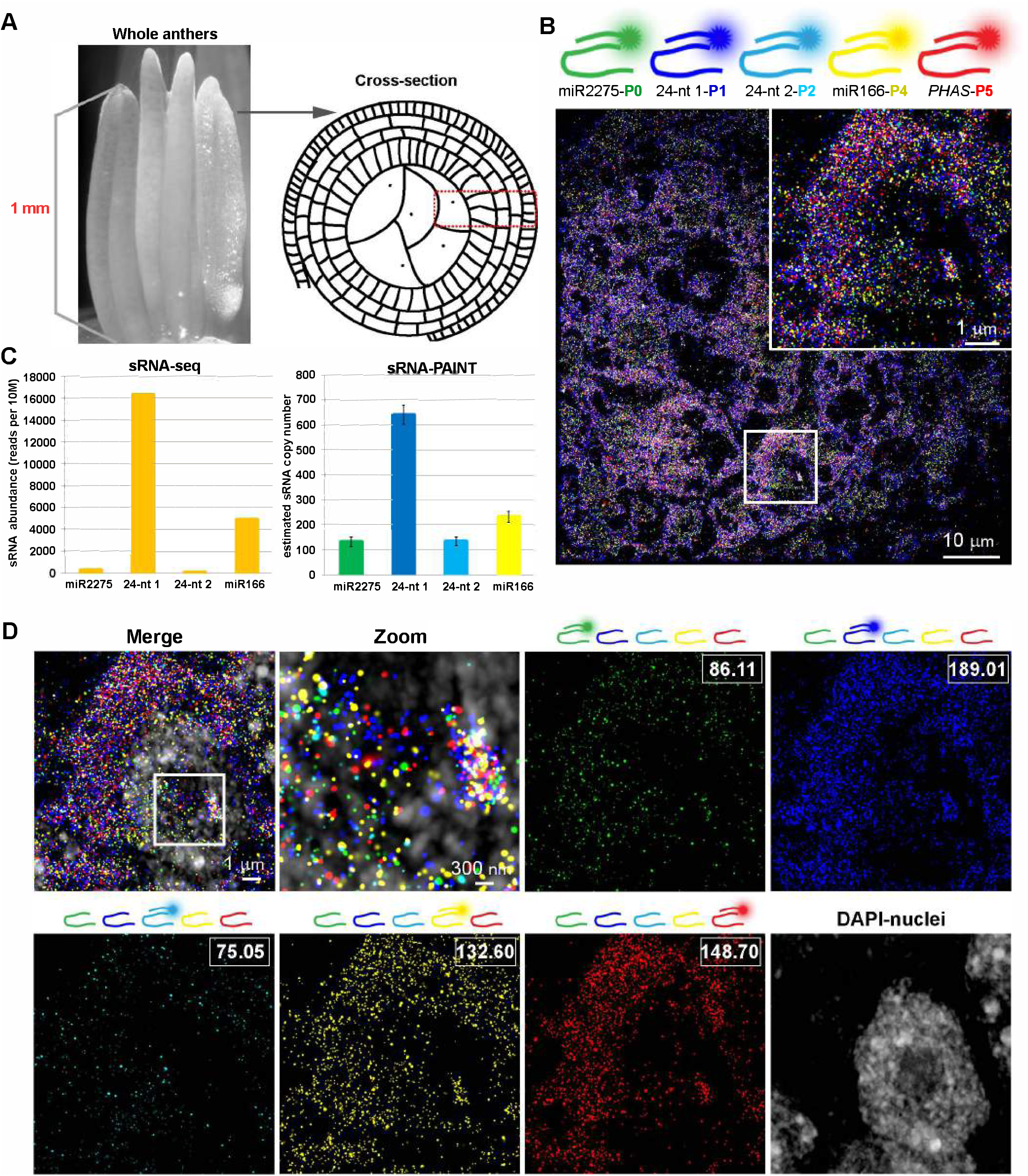
sRNA-Exchange-PAINT for detection of phasiRNA biogenesis components in premeiotic maize anther. (**A**) Schematic of maize anther stage and comparison of material used in sRNA-seq and sRNA-PAINT method. (**B**) Image of sRNA-Exchange-PAINT carried on five targets. Zoomed-in image of the boxed area is shown on the right corner. miR2275 probes were linked with the P0 docking strand and represented with green color. 24-nt phasiRNA 1 probe was linked with P1 docking strand and represented with blue color. 24-nt phasiRNA 2 probe was linked with P2 docking strand and represented with cyan color. miR166 probe was linked with P4 docking strand and represented with yellow color. *PHAS* lncRNA was linked with P5 docking strand and represented with red color. (**C**) Comparison of the quantification result of four target miRNAs using sRNA-seq and sRNA-qPAINT. Representative sRNA copy number estimates were calculated by the sum of the binding sites shown in the boxed area using sRNA-qPAINT. (**D**) Sequential detection of five targets and overlay with the nucleus. Overlay of the channels shows the complexity and density of the small RNAs. Zoomed in image of the boxed area shows single-molecule detection capability with higher-level of multiplexing. The imager strand in each channel is indicated with the diagram above each image, and the quantification of binding sites of each channel is shown in the white box.

## Discussion

sRNA-PAINT provides a robust method that quantitatively detects small RNAs with single-molecule resolution. It greatly improves the resolution and precision of sRNA localization compared to other FISH methods, namely those that are limited by the diffraction limit of light. The small size of sRNAs are generally only detected by one probe per sRNA, as a result, each probe plays an important role with respect to quantification. Our online VARNISH tool assists with the critical probe design step to generate LNA probes with docking strands. Quantification of binding sites then can be deduced from the known binding kinetics of the imager strands to the docking strands with sRNA-PAINT (22). A clear benefit of sRNA-PAINT is ability to multiplex detection beyond three sRNAs, a capability that was previously reserved for longer RNAs.

To develop a quantitative, super-resolution method for sRNA, we chose to design a PAINT-based method over a dSTORM-based method for several reasons. The predictable binding of imager strands to docking strands with PAINT-based method is preferred over the stochastic blinking of dye molecules in dSTORM (22). Furthermore, quantification with a dSTORM probe using a single fluorophore molecule would be greatly hindered by photobleaching. Bleached dye molecules would result in an underestimation of the number of sRNA targets (20, 33). In contrast, qPAINT-based methods are mostly unaffected by fluorophore photobleaching because fluorophore molecules are constantly being replenished by the binding of new imager strands (23). Rather, the major pitfall related to photobleaching is photodamage of the docking strands during qPAINT-based methods caused by the generation of reactive oxygen species (34). After EDC treatment, the photobleaching effect was negligible for the highly abundant 24-nt phasiRNA, since there was no significant difference in the qPAINT quantification after washing and reimaging (Figure 3). EDC has previously been shown to be effective in the stabilization of redox status in treated mammalian tissue (35, 36). Nonetheless, it should be considered for future applications of sRNA-PAINT and it may be beneficial to use a reactive oxygen scavenger buffer during image acquisition (34). Another possibility would be to combine sRNA-PAINT with localization-based fluorescence correlation spectroscopy (lbFCS), rather than qPAINT, for absolute molecular counting of sRNA (37).

An unforeseen benefit of using sRNA-PAINT is that it takes less time than regular *in situ* hybridization methods. sRNA-PAINT samples are directly imaged after hybridization and washing, bringing the sample preparation time down from 4 days to overnight (20). The rapidity was achieved via bypassing the antibody incubation and amplification steps that typify those methods, including our previously reported sRNA-FISH method (20). In sRNA-PAINT, the signal amplification occurs at the same time as image acquisition; although, the drawback is that image acquisition for sRNA-PAINT takes longer than traditional sRNA-FISH methods due to the need to take thousands of images for one field of view for detection. Another benefit of bypassing the antibody amplification steps of traditional *in situ* techniques is that the primary and secondary antibodies add significant amounts of bulk, decreasing the precision of localization by 24 to 30 nm (38). Similarly, the original DNA-PAINT method utilized docking strands conjugated to antibodies for protein detection (22, 23, 39). Overall, the sRNA-PAINT method requires fewer steps than most comparable approaches and is relatively simple. The primary limitation to sRNA-PAINT is the cost of the LNA oligos, which makes it expensive for new probes, although this cost is amortized if the probe is used for many samples, as a typical yield for a purchased LNA oligo is enough for hundreds to thousands of sRNA-PAINT experiments.

Colocalization data from sRNA-PAINT should be evaluated in the context of spatial information at the tissue, subcellular, and molecular levels. In the case of miR2275 and the 24-nt phasiRNA, sRNA-PAINT confirmed *in situ* hybridization data that showed these two sRNAs co-localize at the tissue level in tapetal layer and pollen mother cells, and, at the subcellular level, mainly in the cytoplasm (7, 20). However, at the molecular level, we observed minimal colocalization – evident because sRNA-PAINT has single molecule resolution. The size of a small RNA is about 10 nm (40) and the imaging resolution of our sRNA-PAINT is sub-20 nm. Co-localization of miR2275 and phasiRNA it triggered at the molecular level would suggest that they physically interact with each other or that their density is too high for the method to resolve individual target sRNA molecules; neither of which is apparently the case. In general, colocalization at the single molecule level is plagued by potential imaging artifacts, such as sample drift and sample distortion of deparaffinized sections during repetitive wash and imaging steps. We used the gold fiducials immobilized on coverglasses for lateral image registration and autofocus to correct axial drift; however, these approaches will not solve issues caused by sample damage and distortion over time. Our plans include overcoming these limitations, mainly by mitigating sample distortion, so that colocalization analysis of sRNA-PAINT can be used for sRNA-mRNA (messenger RNA) and sRNA-protein interactions to examine outstanding questions in sRNA biogenesis and processing.

sRNA-PAINT was specifically developed for sRNA, which are too small to apply methods like smFISH that are widely applied for longer mRNAs or lncRNAs. However, here we show that sRNA-PAINT can be used for longer RNAs, but with some caveats. Compared to smFISH, the sRNA-PAINT method detected twice as many copies of *PHAS* lncRNA. Qualitatively, sRNA-PAINT detected binding sites in areas that appeared as diffuse fluorescence by smFISH. We hypothesize that sRNA-PAINT detected both the full length *PHAS* lncRNA and the 5’cleaved products (Figure 4A). These degradation products will have fewer binding sites for smFISH probes and may not result in discrete, bright spots. Therefore, we concede that smFISH is a superior method for quantifying full-length, longer RNAs and should be used for that application. sRNA-PAINT is best suited for multiplexing the detection of longer RNAs with sRNAs or to study the processing of longer RNAs. Improvements in colocalization described above, combined with multiplexing, may lead to powerful advancements in the sRNA-PAINT approach to study sRNA biogenesis from longer precursors. Indeed, the limited number of experiments conducted to develop sRNA-PAINT has led to new hypotheses. For example, we detected a higher number of binding sites for *PHAS* lncRNA than its derivative 24 nt phasiRNA 2. One possibility is the probes detecting mature phasiRNA-2 failed to detect the *PHAS* lncRNA precursor, as that region of the precursor may be predominantly present as dsRNA. As suggested above, another possibility is that the cleaved 5’ portion of the *PHAS* precursor accumulates in the cells after AGO1-directed cleavage (Figure 4).

A distinct advantage of sRNA-PAINT is that it is compatible with Exchange-PAINT, for multiplexing using multiple docking strands that are linked to the same fluorophore (23). Our previously described sRNA-FISH method (20) is limited to only two to three targets, mainly due to the limited choices of antibodies and fluorophores. Achieving the theoretical level of multiplexing for protein detection is mostly limited by the number of primary and secondary antibodies. Our antibody-free method for sRNAs potentially can achieve the theoretical level of multiplexing by designing hundreds of sRNA-PAINT VARNISH probes to query numerous targets in the same sample. The current barriers of reaching the full potential of sRNA-PAINT are the cost of LNA probes and the imaging time. These barriers may soon be removed as the cost of LNA probes has been decreasing and new methods for increasing the speed of PAINT-based methods have been developed (39, 41). However, sRNA-PAINT, as currently conceived, will not reach the same level of other highly multiplexed barcoding methods for mRNAs, such as MERFISH and seqFISH+. MERFISH enabled multiplexed detection of 100 to 1000 RNA species in a single cell using a combination of encoding probes and readout probes (42). The seqFISH+ method was able to detect 10,000 transcripts with high accuracy (43). Our main limitation is the small size of sRNAs. They are simply too small to barcode with non-LNA probes. LNA probes are required for sRNA-PAINT (Figure 2), and high-level multiplexing may be hindered by their cost and the required higher hybridization (55 °C) and stripping (90 °C) temperatures compared to DNA probes. One intriguing avenue to increase multiplexing is to barcode the docking strands. Agasti et al. (44) tested the orthogonality of 52 docking sequences and concluded that those docking sequences could be used as DNA-barcoded labeling probes for PAINT. We anticipate that many advances in our sRNA-PAINT method will be driven by clever, new ways to improve all PAINT-based methods by other research groups.

Finally, our sRNA-PAINT method provides an alternative to conventional sRNA-FISH, as it is capable of higher resolution, quantification, and multiplexing. All FISH-based methods are dependent on the actual hybridization efficiency of the probes prior to imaging. sRNA-PAINT method still retains the pros and cons of any LNA-based in situ method and that PAINT technology quantification is restricted to the number of binding sites (hybridized probes). The development of LNA technology increased the specificity and efficiency of sRNA-FISH methods making the development of sRNA-PAINT possible. Future improvements in probes, such as the use of next-gen bridged nucleic acids (BNAs) (45) may provide more efficient and specific labeling of target sRNAs that will benefit all FISH-based RNA detection methods, including sRNA-PAINT. A drawback of the sRNA-PAINT method is that it can only be applied to fixed tissues or cells. Live-cell imaging, especially aptamer-based sensor imaging for small RNAs, was made possible by fluorescent RNA aptamers such as Spinach and Mango (46-48); however, these methods have low sensitivity and resolution. Creating a live-cell sRNA-PAINT approach with single molecule sensitivity would shed light on the dynamics and ever-changing contents of cellular RNAs.

## Acknowledgements

This project was supported by the US NSF Plant Genome Research Program, awards 1649424, 1611853 and 1754097. We thank members of the Meyers and Caplan labs for help and support. We thank Virginia Walbot (Stanford University) and members of her lab for supplying maize materials and for useful discussions on another small RNAs. We thank Prof. Dr. Ralf Jungmann for his support in DNA-PAINT, qPAINT and analysis with Picasso. Microscopy equipment was acquired with a shared instrumentation grant (S10 OD016361) and access was supported by the NIH-NIGMS (P20 GM103446), the NSF (IIA-1301765) and the State of Delaware.

## Supplementary Figure Legends

**Supplementary Figure 1.**
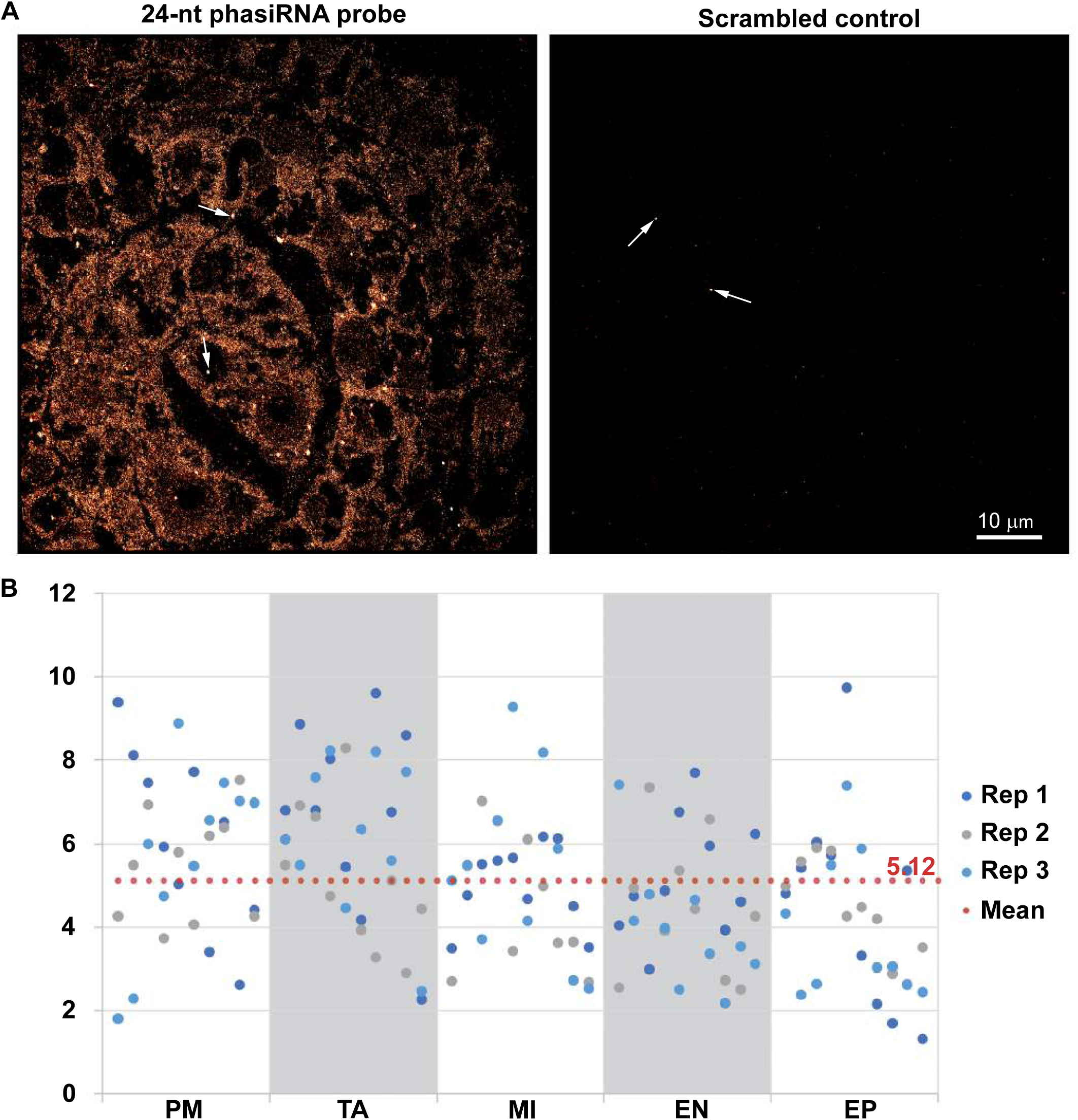
sRNA-PAINT showed limited to no background with a scrambled control probe. (**A**) sRNA-PAINT with probes for a 24-nt phasiRNA and a scrambled control. Both docking strands were detected with corresponding imager strands. Arrows indicate 2 out of ∼66 fiducials used in experiments. Scrambled control detected little or no signal. Scale bar = 10 µm for all images. (**B**) qPAINT analysis of background level of LNA probes with scrambled controls. Number of the binding events for each calculated area (20 pixel size diameter circle in Picasso software) in different cell layers were plotted. A mean of 5.12 binding sites were calculated and used as background level for all subsequent calculations. Ten locations in each cell layer were picked for qPAINT quantification. Three replicates and a total of 150 samples were used for the background calculation (n=150).

**Supplementary Figure 2.**
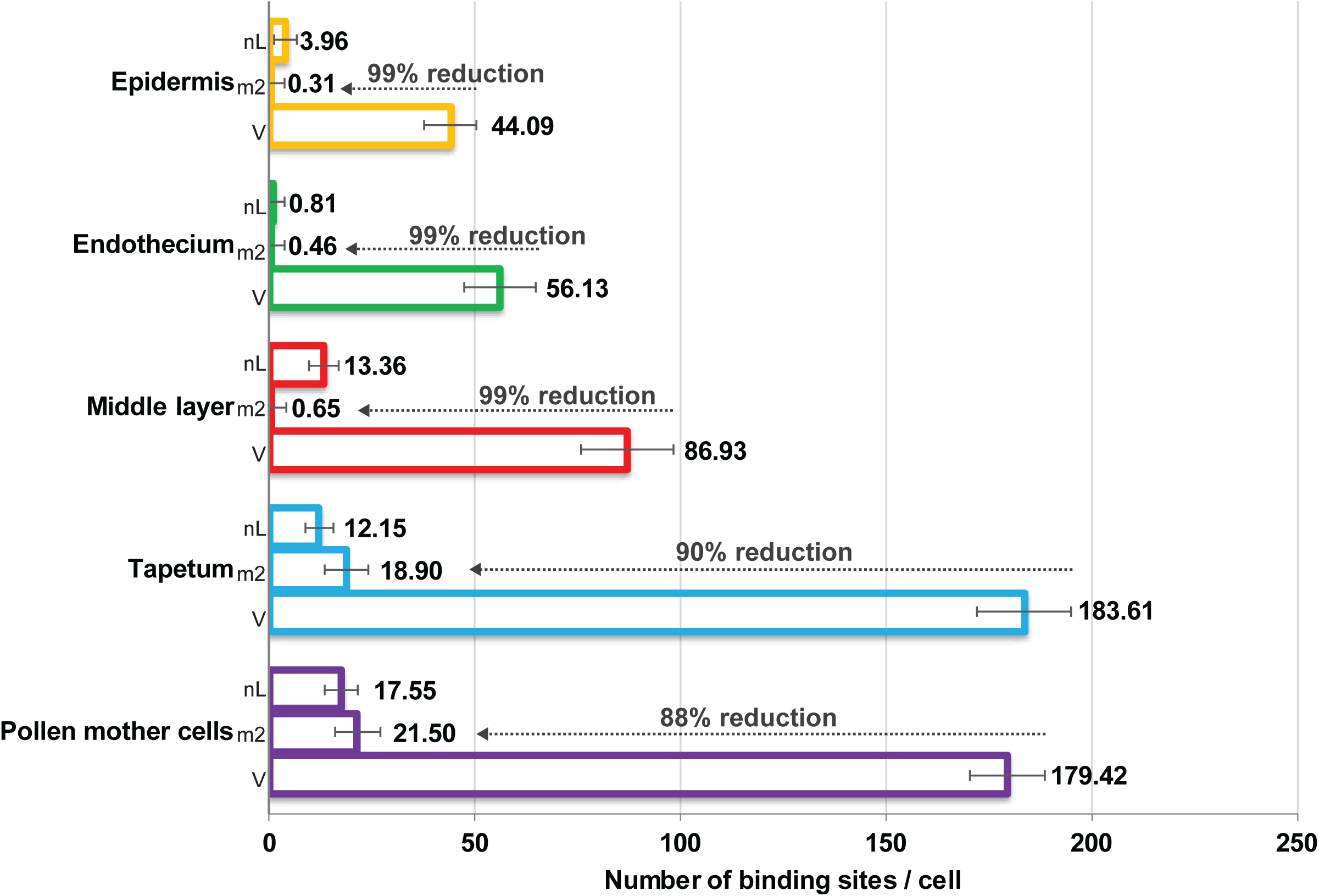
Quantification of specificity of VARNISH probes. VARNISH 24-nt phasiRNA probe (V), a probe with 2-nt mutations (m2), and probe no LNAs (nL) were applied to each anther sample. Quantification was done using qPAINT analysis. Numbers show the binding events for each cell in different cell layers. An average of 95% reduction is detected with m2 in all cell layers. 10 different locations in each specified cell layer and three replicates (two replicates for nL) were taken for statistical analysis (n=150 for V and m2, n=100 for nL). A no LNA probe with 2 nt mutation was used as control for nL probe for qPAINT calculation. Letters indicate significant differences using Tukey’s test (P < 0.05).

**Supplementary Figure 3.**
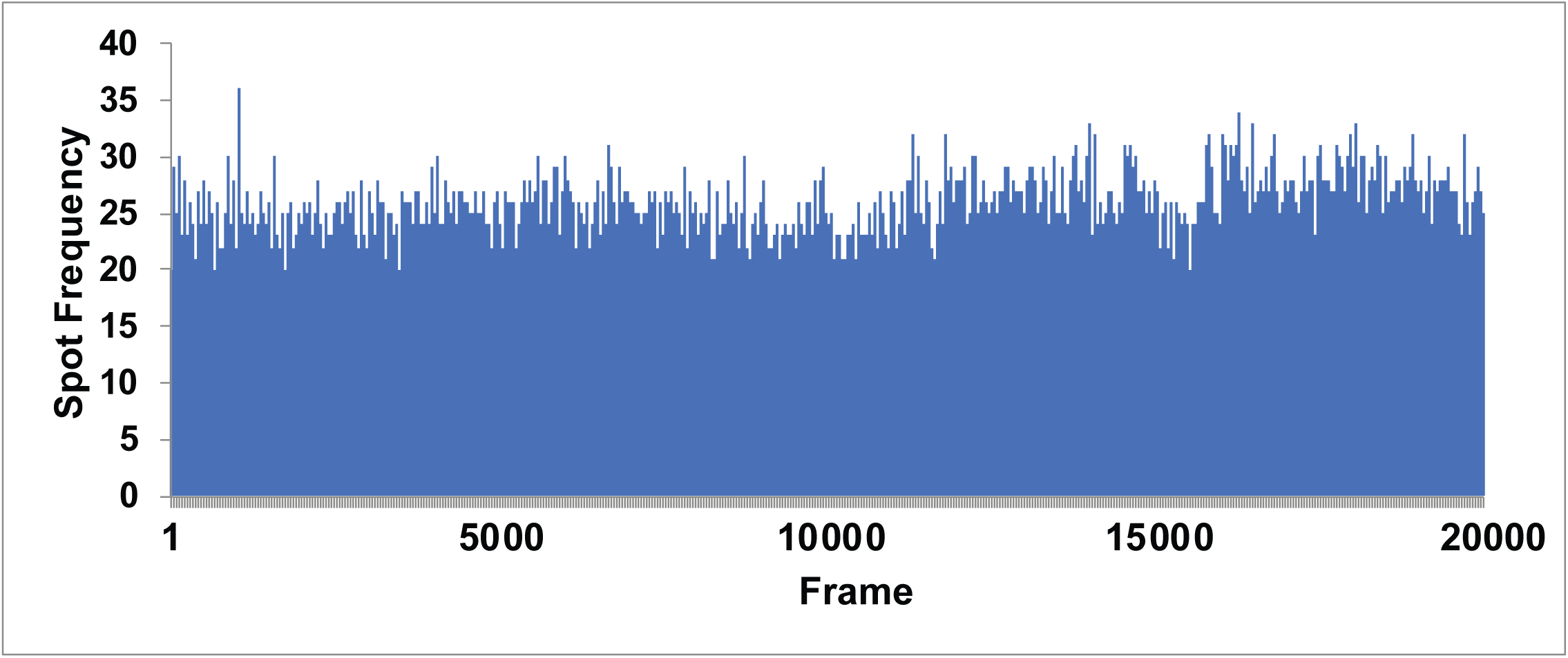
VARNISH probe signals remain stable during imaging. Number of localization spots detected remain relatively stable during the imaging process of 20,000 frames for 24-nt phasiRNA.

**Supplementary Figure 4.**
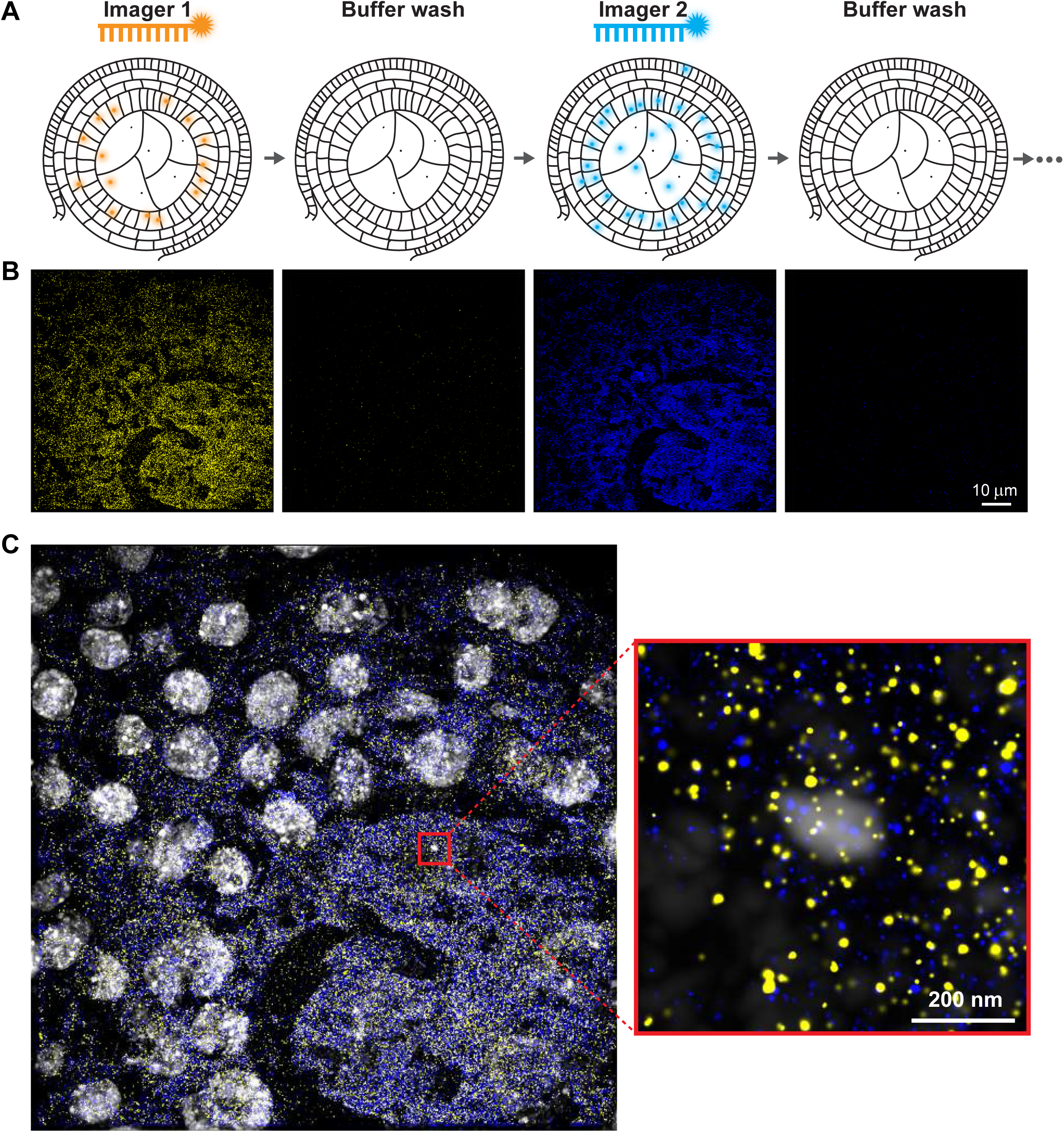
sRNA-Exchange-PAINT for miR2275 and 24-nt phasiRNA co-localization. (**A**) A diagram showing sRNA-Exchange-PAINT, which differs from sRNA-PAINT via the use of multiple imager strands. Imager strand 1 is first introduced to the sample, and after imaging, imager strand 1 is removed with a buffer wash. Next, imager strand 2 is introduced. This process can be repeated until the needed number of targets is reached. (**B**) sRNA-Exchange-PAINT on miR2275 (yellow) and a 24-nt phasiRNA (blue) in maize anther. After the buffer wash, imaging showed little or no signal following each round of detection using the imager strands. Scale bar = 10 µm for all images. (**C**) Merged and zoomed image of the marked region is shown. These two sRNAs do not show significant co-localization at nanometer resolution.

## Supplementary Tables

*Supplementary Table 1. Information for sequential image of phasiRNA generation components using sRNA-Exchange-PAINT*.

*Supplementary Table 2. Probes for smFISH of a* PHAS *precursor*.

## Notes

### Competing Interest Statement

The authors have declared no competing interest.

### Summary of Updates

We repeated nearly all experiments using EDC to prevent probe loss during wash steps. We calculated background binding of a scramble LNA probe and used this to more accurately estimate copy numbers and corrected a quantification mistake. We also introduced two mutations to our miR2275 probe and show an average 95% reduction in binding site detection. Lastly, we conducted the challenging experiment of sRNA-PAINT and smFISH on the same sample.

